# Cold Adaptation Leads To Repolarization Gradients Development In The Rainbow Trout Heart

**DOI:** 10.1101/2021.03.22.436433

**Authors:** Marina A. Vaykshnorayte, Vladimir A. Vityazev, Jan E. Azarov

## Abstract

**Introduction:** Thermal adaptation in fish is accompanied by morphological and electrophysiological changes in the myocardium. Little is known regarding changes of spatiotemporal organization of ventricular excitation and repolarization processes with acclimatization. We aimed to evaluate transmural and apicobasal heterogeneity of depolarization and repolarization characteristics in the in-situ heart of rainbow trout in seasonal acclimatization.

**Methods:** The experiments were done in the summer-acclimatized (SA, 18°C, n=8) and winter-acclimatized (WA, 3°C, n=8) rainbow trout. 24 unipolar electrograms were recorded with plunge needle electrodes (eight lead terminals each) impaled into the ventricular wall. Activation time (AT), end of repolarization time (RT), and activation-repolarization interval (ARI, a surrogate for action potential duration) were determined as dV/dt min during QRS-complex, dV/dt max during T-wave, and RT-AT difference, respectively.

**Results:** The SA fish demonstrated relatively flat apicobasal and transmural AT and especially ARI profiles. In the WA animals, ATs and ARIs were longer as compared to SA animals (p≤0.001), ARIs were shorter in the compact layer than in the spongy layer (p≤0.050), and within the compact layer, the apical region had shorter ATs and longer ARIs as compared to the basal region (p≤0.050). In multiple linear regression analysis, ARI duration was associated with cardiac cycle duration and AT in SA and WA animals. The WA animals demonstrated additionally an independent association of ARIs with spatial localization across the ventricle.

**Conclusion:** Adaptation to cold conditions in rainbow trout was associated with a spatial ventricular remodeling leading to the development of repolarization gradients typically observed in mammalian myocardium.

**SUMMARY STATEMENT:** The study gives an example of thermal adaptation in fish realized at the level of spatiotemporal organization of myocardial depolarization and repolarization.

## INTRODUCTION

Spatiotemporal organization of ventricular excitation in vertebrates is determined by the spread of activation wave(s) and nonuniform repolarization. The former is determined by the morphology of conduction fibers, which sets mainly endo-to-epicardial activation sequence in most vertebrates. The nonuniformities of repolarization manifest as so-called repolarization gradients (differences in action potential durations between ventricular regions). At least, interventricular, apicobasal and transmural repolarization gradients can be discerned in the ventricular myocardium (Arteyeva et al., 2013, Meijborg et al., 2014). The magnitude of these gradients varies significantly across animal species and experimental conditions. Though the transmural gradient is elusive (Boukens et al., 2017), careful measurements usually demonstrate at least the small difference in repolarization timing between subendocardial and subepicardial regions, and the electrophysiological nature of these difference was shown long ago (Antzelevitch et al., 1991).

Ventricular myocardium in fish has spongy and compact muscle layers (Duran et al., 2015, Farrell et al., 2009, Kochova et al., 2015, Pieperhoff et al., 2009) expressed to a different extent in different species. The spongy layer is considered as a predecessor of a His-Purkinje system (Jensen et al., 2012), while the compact layer bears the most contractile load and is related to coronary vasculature (Farrell et al., 2009). It might be expected that the presence of the two different muscle layers would cause the development of the transmural repolarization gradient. However, the previous works did not demonstrate the difference in action potential duration between the inner and outer layers (Vaykshnorayte et al., 2011a, Patrick et al., 2011).

Thermal adaptation in fish is critical for survival, and the function of the heart is essential for this process (Farrell et al., 2009). Thermal adaptation modifies deeply heart morphology and functional characteristics (Keen et al., 2017). Specifically, lowering the ambient temperature leads to a relative decrease of compact myocardium and the development of fibrosis in the ventricular wall, which could result in uncoupling of the compact and spongy layers (Keen et al., 2016, Klaiman et al., 2011). The temperature-dependent changes of cardiac electrophysiological properties that determine heart rate and conduction were shown to play an important role in seasonal acclimatization (Vornanen, 2016). However, the contractile function of the myocardium depends on coordination of the intraventricular electrical properties (Markhasin et al., 2012), but little is known up to date about changes of spatiotemporal patterns of ventricular depolarization and repolarization in thermal adaptation. It might be expected that thermal acclimatization modifies the ventricular electrophysiological patterns on different anatomical axes. In the present study, we tested this hypothesis by evaluating apicobasal and transmural profiles of depolarization and repolarization timing in summer-acclimatized (SA) and winter-acclimatized (WA) rainbow trout.

## MATERIALS AND METHODS

The investigation was carried out in accordance with the *Guide for the Care and Use of Laboratory Animals, 8th Edition* published by the National Academies Press (US) 2011 and was approved by the institutional ethical committee.

8 SA (18°C, August) and 8 WA (3°C, April) rainbow trout (Onchorhynchus mykiss; body weight 300-600 g) were studied at the fishery farm (located in Kirov Region, Russia, 58°40’ N; 50°47’ E). The measurements were performed at an ambient temperature approximately 2°C higher than the temperature of acclimatization. Fish were stunned by a sharp blow at the head and the spine was cut. After that, the heart was exposed via the ventral surface. Ventricular unipolar electrograms were recorded from the ventricular walls as described earlier (Vaykshnorayte et al., 2011a). In brief, three plunge needle electrodes were inserted in the ventricular wall at the apex, anterior part of the base, and in the posterior region adjacent to the atrioventricular junction. Each needle electrode bore eight lead terminals connected to the amplifiers of a custom-designed system for electrophysiological contact mapping (16 bits; bandwidth 0.05 to 1000 Hz; sampling rate 4000 Hz).

In each ventricular lead, activation time (AT) and end of repolarization time (RT) were determined as the instants of dV/dt minimum during QRS complex and dV/dt maximum during T-wave, respectively (Coronel et al., 2006). An activation-repolarization interval (ARI, a surrogate for the action potential duration) was determined as the time differences between the RT and AT. The length of a plunge needle electrode was selected to match the thickness of the ventricular wall. Two outermost and two innermost lead terminals were referred to as compact layer and spongy layer terminals, respectively. In order to get one value for each layer, the data obtained from the two corresponding leads were averaged.

Data are expressed as medians and interquartile intervals. Statistical analysis was performed with SPSS package (IBM SPSS Statistics 23, SPSS, Inc., Chicago, Illinois, USA). Wilcoxon and Fridman tests were applied for paired and multiple comparisons, respectively, within the same groups of animals. Mann-Whitney test was performed for comparisons between the WA and SA animals. The differences were considered significant at p<0.05.

## RESULTS

Thermal adaptation in rainbow trout resulted in significant changes in cardiac electrophysiological properties. Figure 1 displays representative electrograms recorded in different ventricular areas in the subepicardial and subendocardial regions. Both activation and repolarization processes were prolonged in the WA animals. At all studied sites, ATs and ARIs were significantly greater in cold conditions (p<0.001). Cardiac cycle duration was longer in the WA as compared to SA animals [median 4045 (IQR 2360-5230) ms vs 1991 (IQR 1406-2767) ms, p=0.029, respectively].

**Figure 1.**
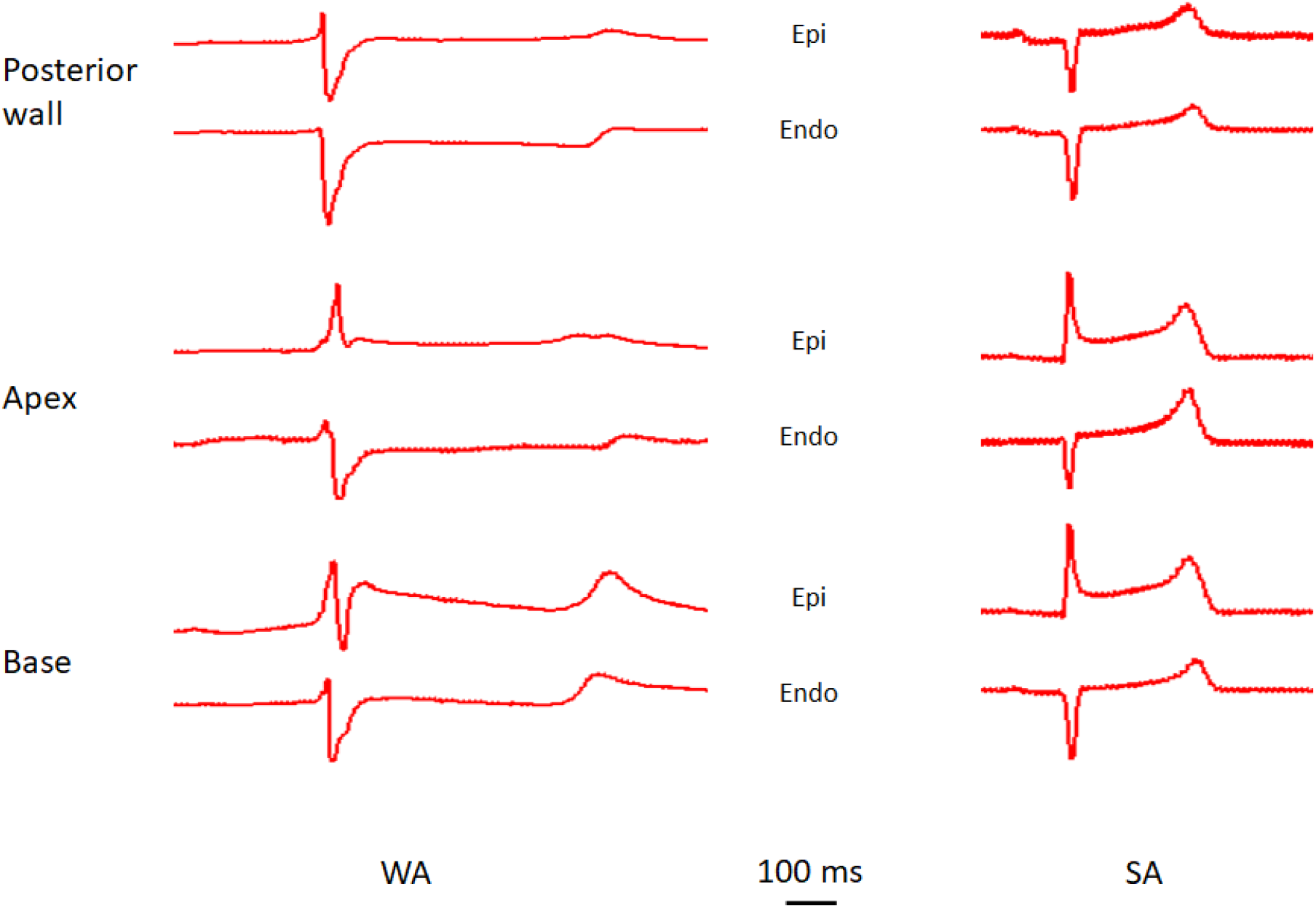
Representative subepicardial (epi) and subendocardial (endo) electrograms from the posterior wall near the atrioventricular junction, apex, and base of the ventricle in the winter acclimatized (WA) and summer acclimatized (SA) rainbow trout. See prolonged QRS and QT intervals in WA fish. The predominant polarity of QRS complexes is negative in the subendocardial layers and positive in the subepicardial layers reflecting endo-to-epi activation spread in the base and apex of both WA and SA fish. In the SA fish, the T-waves were uniform across the ventricular myocardium. In contrast, the WA fish demonstrated different T-wave polarity and/or morphology in different layers and areas of the ventricle reflecting heterogeneous repolarization.

We first analyzed parameters of ventricular depolarization and repolarization in different ventricular areas averaged over wall thickness. An earliest ventricular activation was consistently observed in the posterior wall near the atrioventricular junction in the SA and WA fish. From this region, activation wave spread to the apex and anteriorly resulting in that the ATs in the apex and anterior base were significantly longer as compared to the posterior wall both in SA and WA fish (Figure 2, panel A). Repolarization duration distribution on the apicobasal axis was relatively uniform. The ARI durations did not differ between the three ventricular areas in both groups of animals (Figure 2, panel B).

**Figure 2.**
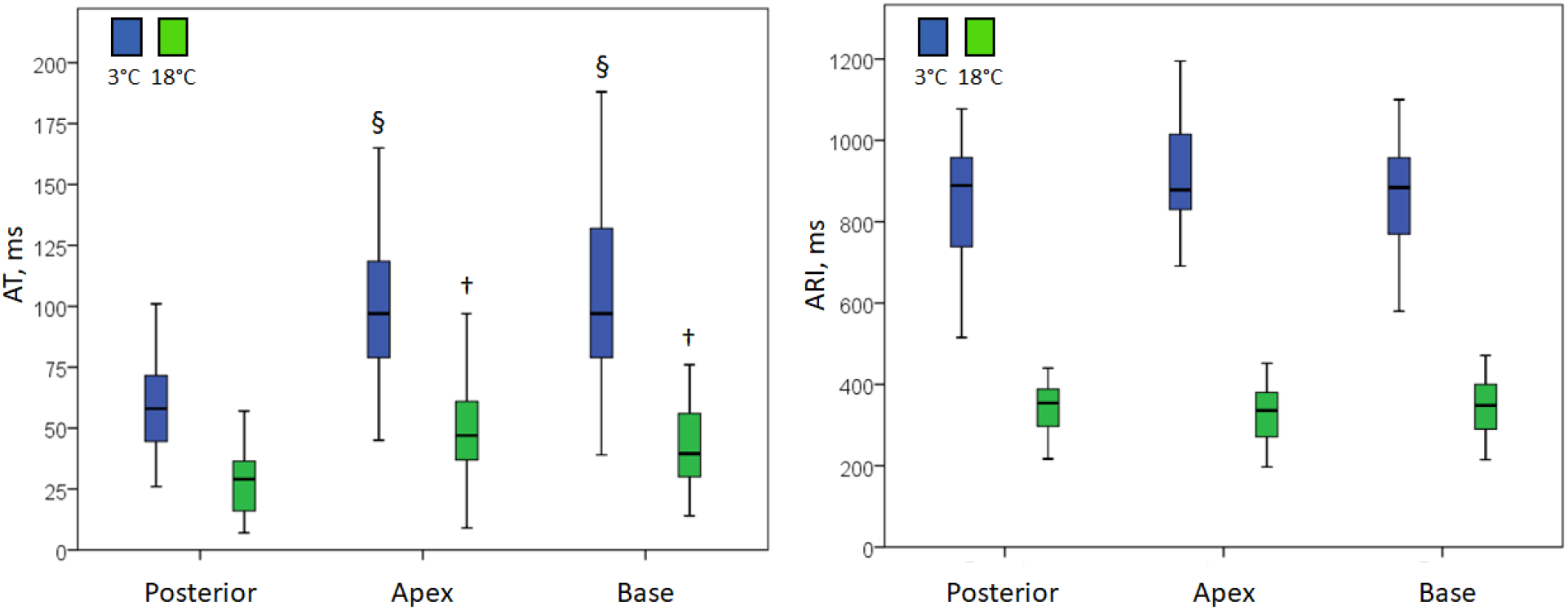
Activation times (AT) and activation-repolarization intervals (ARI) averaged for the entire thickness of the ventricular wall in different areas of the ventricle. See longer temporal parameters in the WA animals (p≤0.001 for all comparisons) and significantly earlier activation of the posterior wall near atrioventricular junction (Posterior) as compared to the heart apex (Apex) and anterior base (Base). See also a relatively flattened ARI distribution across the above areas both in SA and WA animals. † p=0.008 vs Posterior, § p=0.012 vs Posterior.

Then, separately in the compact (subepicardial) and spongy (subendocardial) layers, we compared the timing of depolarization and repolarization between the apex and base (Figure 3). The SA fish had a relatively uniform distribution of both ATs and ARIs in both layers demonstrating that the apical and basal regions are activated and repolarized at approximately the same time. The WA animals had a similar uniform apicobasal distribution of depolarization and repolarization timing in the spongy layer. However, apicobasal differences were observed in the compact layer in the WA fish. Here, the apical region had a shorter AT and longer ARI as compared to the subepicardial basal region.

**Figure 3.**
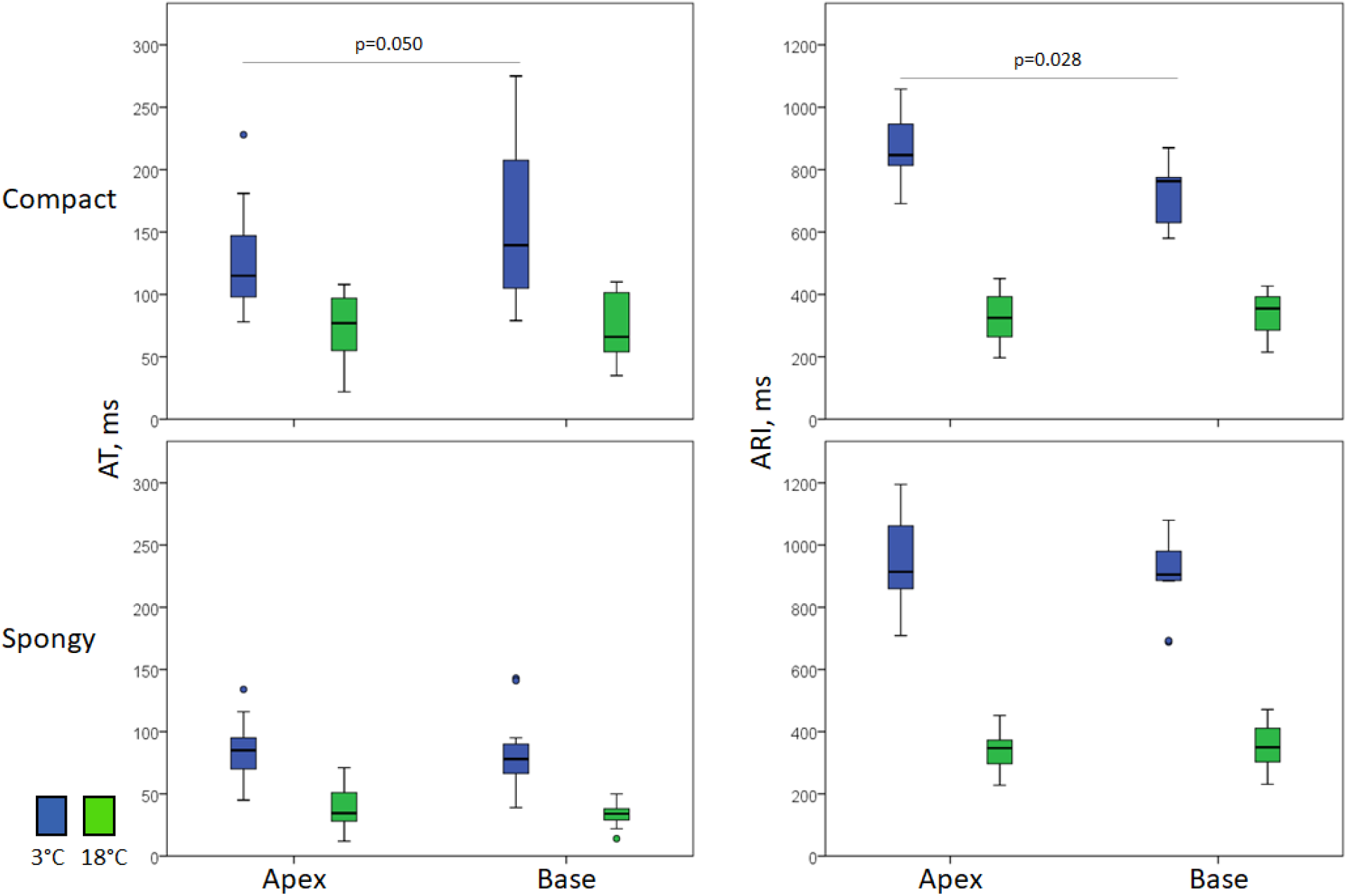
Apicobasal distribution of activation times (AT) and activation-repolarization intervals (ARI) separately in the compact and spongy layers in SA and WA animals. See the opposite distribution of ATs and ARIs in the compact layer of WA animals.

On the transmural axis both in SA and WA rainbow trout, activation wave spreads from endocardium to epicardium. Figure 4 demonstrates that the subendocardial ATs were shorter than the subepicardial ATs at both temperatures of acclimatization. The transmural profile of repolarization duration was relatively flat at 18°C (Figure 5). However, in the WA animals, a significant transmural difference in ARI durations was observed in all studied ventricular areas with the subepicardial ARIs being shorter than the subendocardial ARIs (Figure 5).

**Figure 4.**
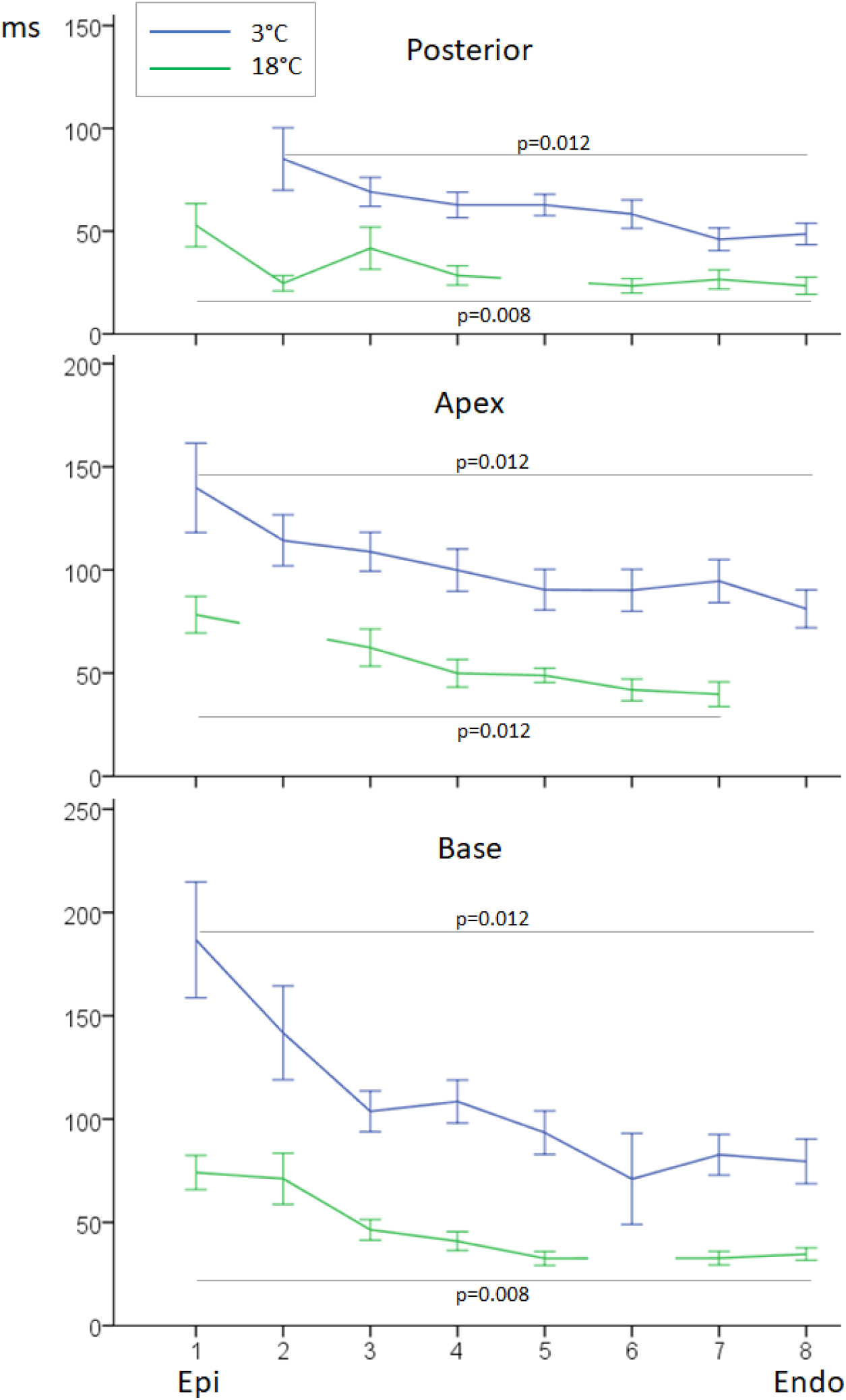
Transmural profiles of activation times (AT) in SA (18°C) and WA (3°C) rainbow trout in the posterior wall near the atrioventricular junction (Posterior), apex (Apex), and anterior base (Base). The numbers from 1 to 8 on the horizontal axis indicate leads on the intramural needle from epicardium (lead 1) to endocardium (lead 8). The missing points relate to bad quality signals from some leads. See the progressive increase of ATs from endocardium to epicardium especially in the WA animals.

**Figure 5.**
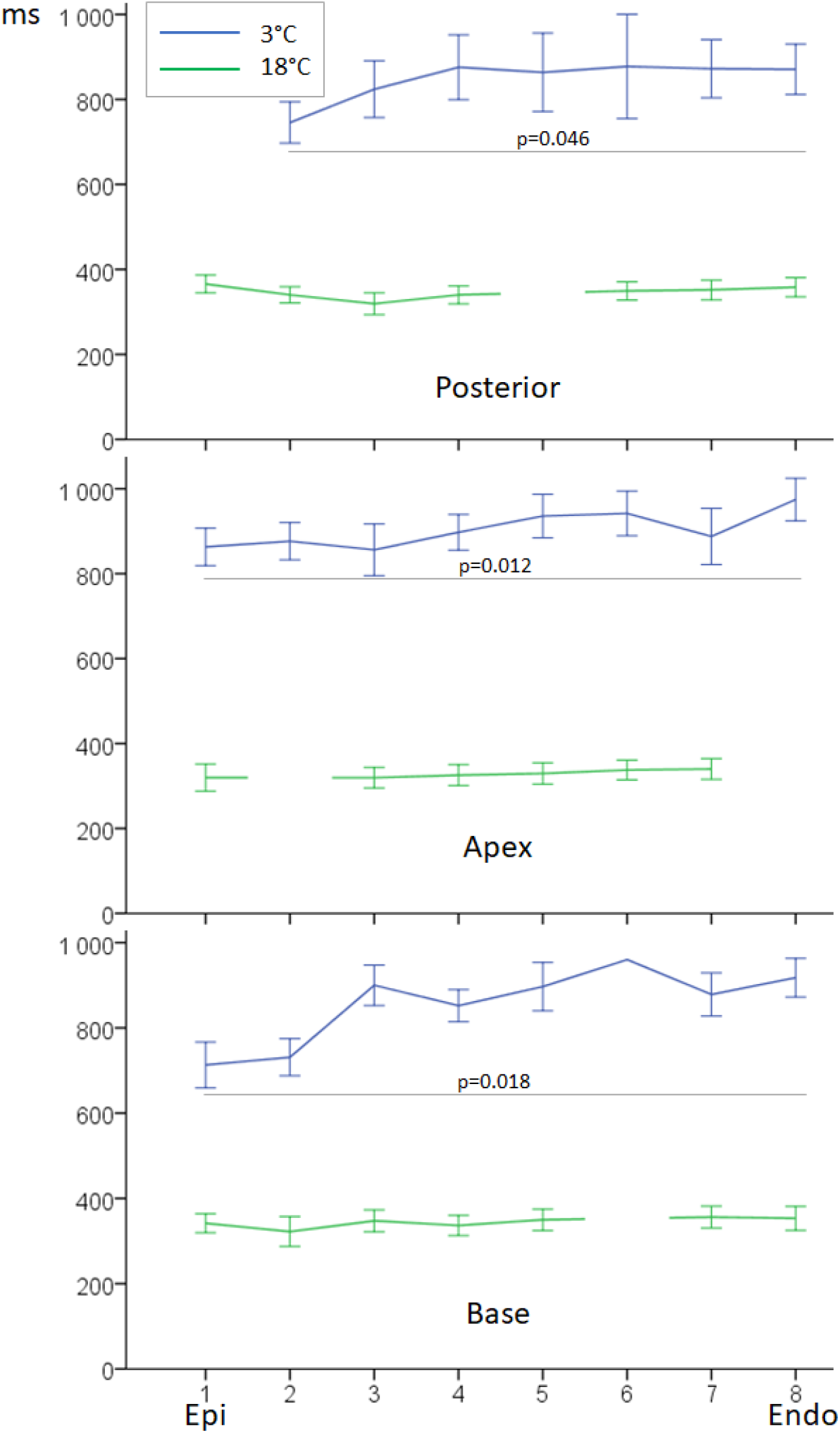
Transmural profiles of activation-repolarization intervals (ARI) in SA (18°C) and WA (3°C) rainbow trout in the posterior wall near the atrioventricular junction (Posterior), apex (Apex), and anterior base (Base). The numbers from 1 to 8 on the horizontal axis indicate leads on the intramural needle from epicardium (lead 1) to endocardium (lead 8). The missing points relate to bad quality signals from some leads. See a flat ARI distribution in the SA fish and progressive increase of ARIs from epicardium to endocardium in the WA fish.

In multivariate regression analysis (Table 1), we tested the association of ARI with the factors expected to affect repolarization duration: a cardiac cycle length (RR interval), AT, and position on the transmural and apicobasal axes. ARI correlated positively with cardiac cycle length and negatively with AT both in the SA and WA animals (Figure 6). However, the ARIs in the WA fish also demonstrated a significant association with spatial positions independently of RR interval and AT (Table 1).

**Table 1.** Associations of ARI durations with electrophysiological and spatial factors in SA (18°C) and WA (3°C) rainbow trout.

| Predictor | SA |  | WA |  |
| --- | --- | --- | --- | --- |
|  | B (95% CI) * | p | B (95% CI) | p |
| RR | 14.77 (9.17 – 20.36) | <0.001 | 64.32 (49.13 – 79.50) | <0.001 |
| AT | -0.95 (-1.40 – -0.49) | <0.001 | -0.82 (-1.51 – -0.14) | 0.019 |
| Transmural position ** | -2.15 (-6.73 – 2.43) | 0.355 | 12.23 (1.57 – 22.89) | 0.025 |
| Apicobasal position *** | 6.35 (-5.17 – 17.88) | 0.278 | 46.53 (14.92 – 78.14) | 0.004 |

\* - regression coefficient and 95% confidence interval, \*\* - ordinal variable from epi to endo; \*\*\* - nominal variable, AT – activation time, RR – RR interval in ECG (cardiac cycle length).

**Figure 6.**
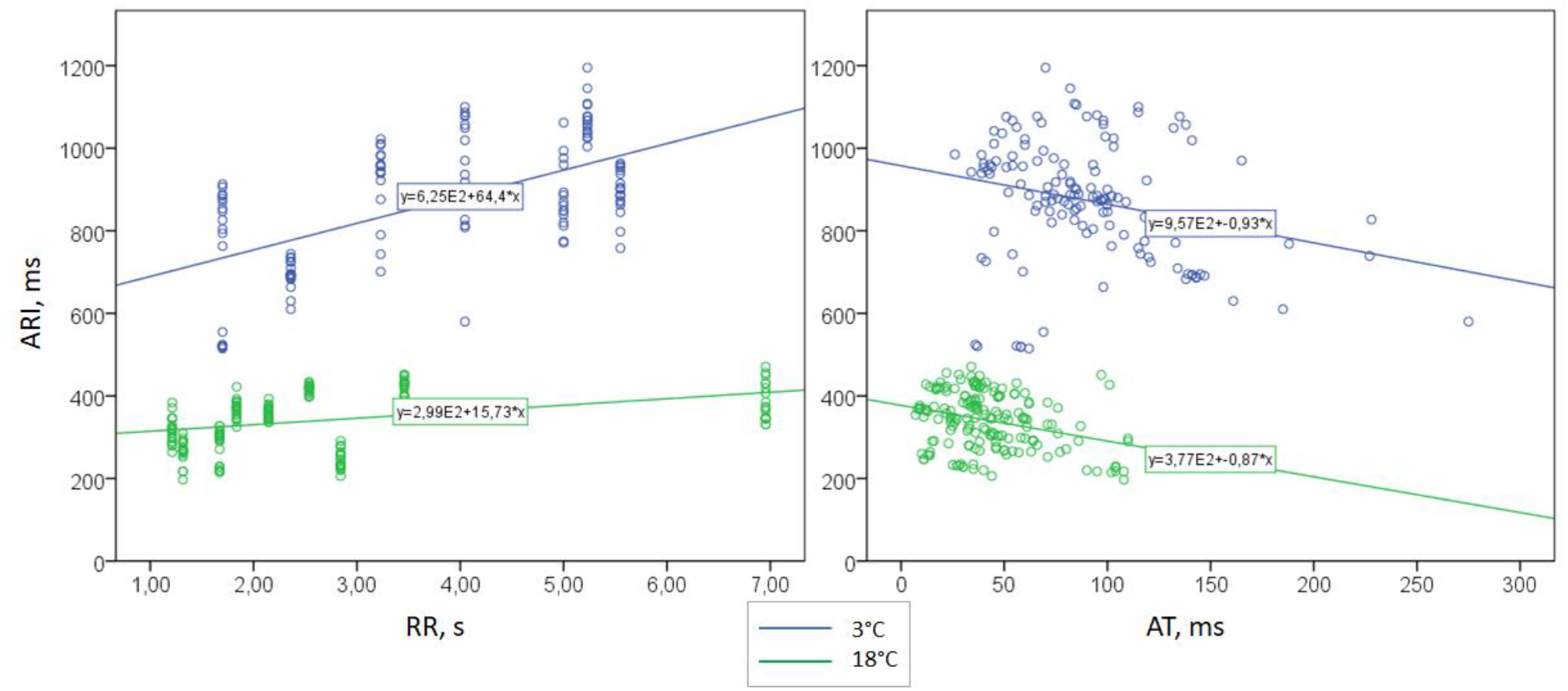
Scatter plots activation-repolarization intervals duration (ARI) vs cardiac cycle length (RR interval) and activation time (AT) with regression equations. See a significant direct relationship between ARI and RR (left) and an inverse relationship between ARI and AT (right).

## DISCUSSION

The present study demonstrated the development of electrophysiological heterogeneities in the rainbow trout heart related to cold adaptation. It was shown that the transmural repolarization duration gradient from epicardium to endocardium (opposite to the depolarization gradient) formed in the WA animals. Furthermore, cold acclimatization was associated with the development of the opposite apicobasal activation and repolarization duration gradients in the compact layer of the ventricle.

Cold conditions led to slowing electrophysiological processes, which manifested as the decreased heart rate (prolonged cardiac cycle length), prolonged ventricular activation (delayed ATs), and repolarization (prolonged ARIs). Prolonged ventricular activation is likely due to the reported accumulation of connective tissue in the ventricular wall caused by cold acclimation (Keen et al., 2016, Klaiman et al., 2011). Prolongation of repolarization and heart rate decrease was observed despite the expected temperature compensation (Haverinen and Vornanen, 2007, Aho and Vornanen, 2001, Hassinen et al., 2008, Abramochkin and Vornanen, 2015). These observations could be ascribed to a combination of temperature compensation and direct thermal effects. Moreover, the heart rate slowing in WA animals might be related to the elimination of extracardiac regulation due to the damage of the central nervous system. The mechanism of ARI prolongation should be based on the altered balance between depolarizing and repolarizing currents in favor of depolarization. The most plausible explanation for these cold-induced changes of ARI should be thermal suppression of IKr (for review, see (Vornanen, 2016)).

Probably, the most important finding of the present study is that repolarization prolonged heterogeneously in cold conditions with the development of repolarization duration gradients opposite to activation sequence on the corresponding directions, namely ARIs were longer in the apex than in the base within the external (compact) layer of the myocardium, and progressively increased from epicardium to endocardium on the transmural axis. Previously, our group reported that acute heart cooling induced predominant prolongation of repolarization in the apical region of the heart both on the epicardial surface of amphibian (Vaĭkshnoratĭe et al., 2007) and mammalian (Azarov et al., 2008) heart. In the present study, we found that a similar change developed in seasonal acclimatization in fish heart and that this effect is limited to the compact myocardial layer.

This study is the first, to our knowledge, to report the presence of a transmural direction of activation spread in the fish heart. In contrast to the previous work in pikes (Vaykshnorayte et al., 2011a), it is here shown in the rainbow trout ventricle that the activation wave propagated not only along the walls but also from endocardium to epicardium in a similar way as in other vertebrates. This activation pattern was found both in SA and WA fish. In mammals, the transmural gradient of repolarization durations has been also recorded (Arteyeva et al., 2013, Meijborg et al., 2014); however, it is noteworthy that it is much easily found in isolated myocardial preparations than in a whole heart (Boukens et al., 2017). What was shown in the present study is that the transmural repolarization duration gradient can be also observed in the fish ventricle and this gradient is subject to thermal adaptation since it was demonstrated only in WA but not SA animals.

The differences in repolarization durations observed across ventricular myocardium can be due to regional intrinsic electrophysiological differences (expression of channel proteins etc) and electrotonic interaction between adjacent sites (Vaykshnorayte et al., 2011b). The latter results in that the closely spaced myocardial regions tend to finish the repolarization process at the same time. That means that the earlier activated regions would have longer action potential durations, while those activated later would have shorter action potential durations. In other words, electrotonic interaction would result in the inverse relationship between ATs and ARIs with no prerequisites of any intrinsic spatial differences in ion transport machinery. Importantly, such a mechanism suggests that the differences in repolarization durations between the regions should be comparable with corresponding differences in activation times. However, in some cases, the observed difference in repolarization durations is far greater than the AT difference. This situation suggests that the nonuniformities of repolarization could not be entirely ascribed to the electrotonic interaction and there are some intrinsic differences between the regions in question, which could develop during ontogeny or form due to remodeling processes.

To find out whether ARI distribution in WA and SA rainbow trout was governed just by electrotonic interaction or any seasonal remodeling process is involved, we performed multivariate regression analysis (Table 1 and Figure 6). As tested predictors, it included cardiac cycle length, AT, and a spatial location on the transmural and apicobasal axes. If the spatial location is independently associated with ARI duration, it suggests the presence of intrinsic repolarization heterogeneities. We found such an association in the WA but not SA animals that could be explained by myocardial electrical remodeling during cold adaptation resulting in the development of transmural and apicobasal heterogeneities of repolarization durations.

The obtained data suggest that cold adaptation-associated remodeling is related to the increased hemodynamical load in cold conditions. At least part of the observed WA vs SA differences was related to the compact myocardial layer that is considered as a predecessor of contractile myocardium (Jensen et al., 2012). Moreover, apicobasal as well as transmural repolarization distribution in the WA fish is spatially inversely related to the activation sequence, i.e. the earlier the AT, the longer the ARI, and vice versa. This activation-repolarization pattern can facilitate the contraction process, since optimal coordination of the contractile elements requires that activation and therefore contraction proceeds from “slow” to “fast” contracting muscle elements that have long and short action potential duration, respectively (Markhasin et al., 2012).

## CONCLUSION

Thus, the present study demonstrated that thermal adaptation can be based not only on changes in cardiac structure and heart rate but also can involve electrical remodeling of ventricular myocardium leading to a spatial redistribution of repolarization durations. It warrants further investigation of the mechanism and functional significance of found seasonal electrical remodeling.

## LIST OF SYMBOLS AND ABBREVIATIONS

ARI: activation-repolarization interval
AT: activation time
RT: repolarization time
SA: summer acclimatized
WA: winter acclimatized

## ACKNOWLEDGEMENTS

The authors highly appreciate the excellent assistance of Dmitry Bayev, Oleg Alypov and Yuriy Zevakin, the management and staff of Neptun Ltd (Isakovtsy, Kirov Region, Russia), in rearing fish and help during experiments.

## AUTHOR CONTRIBUTIONS

Concept – JEA, design and experiments – MAV, VAV, JEA, data analysis and draft writing – MAV and JEA, critical revision – VAV.

## DATA AVAILABILITY

The data underlying the present research can be requested from the authors.

## REFERENCES

Abramochkin, D. V. & Vornanen, M. 2015. Seasonal acclimatization of the cardiac potassium currents (IK1 and IKr) in an arctic marine teleost, the navaga cod (Eleginus navaga). J Comp Physiol B, 185, 883–90.

Aho, E. & Vornanen, M. 2001. Cold acclimation increases basal heart rate but decreases its thermal tolerance in rainbow trout (Oncorhynchus mykiss). J Comp Physiol B, 171, 173–9.

Antzelevitch, C., Sicouri, S., Litovsky, S. H., Lukas, A., Krishnan, S. C., Di Diego, J. M., Gintant, G. A. & Liu, D. W. 1991. Heterogeneity within the ventricular wall. Electrophysiology and pharmacology of epicardial, endocardial, and M cells. Circ Res, 69, 1427–49.

Arteyeva, N. V., Goshka, S. L., Sedova, K. A., Bernikova, O. G. & Azarov, J. E. 2013. What does the T(peak)-T(end) interval reflect? An experimental and model study. J Electrocardiol, 46, 296.e1-8.

Azarov, J. E., Shmakov, D. N., Vityazev, V. A., Roshchevskaya, I. M., Arteyeva, N. V., Kharin, S. N. & Roshchevsky, M. P. 2008. Ventricular repolarization pattern under heart cooling in the rabbit. Acta Physiol (Oxf), 193, 129–38.

Boukens, B. J., Meijborg, V. M. F., Belterman, C. N., Opthof, T., Janse, M. J., Schuessler, R. B., Coronel, R. & Efimov, I. R. 2017. Local transmural action potential gradients are absent in the isolated, intact dog heart but present in the corresponding coronary-perfused wedge. Physiol Rep, 5.

Coronel, R., De Bakker, J. M. T., Wilms-Schopman, F. J. G., Opthof, T., Linnenbank, A. C., Belterman, C. N. & Janse, M. J. 2006. Monophasic action potentials and activation recovery intervals as measures of ventricular action potential duration: Experimental evidence to resolve some controversies. Heart Rhythm, 3, 1043–1050.

Duran, A. C., Lopez-Unzu, M. A., Rodriguez, C., Fernandez, B., Lorenzale, M., Linares, A., Salmeron, F. & Sans-Coma, V. 2015. Structure and vascularization of the ventricular myocardium in Holocephali: their evolutionary significance. J Anat, 226, 501–10.

Farrell, A. P., Eliason, E. J., Sandblom, E. & Clark, T. D. 2009. Fish cardiorespiratory physiology in an era of climate changeThe present review is one of a series of occasional review articles that have been invited by the Editors and will feature the broad range of disciplines and expertise represented in our Editorial Advisory Board. Canadian Journal of Zoology, 87, 835–851.

Hassinen, M., Haverinen, J. & Vornanen, M. 2008. Electrophysiological properties and expression of the delayed rectifier potassium (ERG) channels in the heart of thermally acclimated rainbow trout. Am J Physiol Regul Integr Comp Physiol, 295, R297–308.

Haverinen, J. & Vornanen, M. 2007. Temperature acclimation modifies sinoatrial pacemaker mechanism of the rainbow trout heart. Am J Physiol Regul Integr Comp Physiol, 292, R1023–32.

Jensen, B., Boukens, B. J., Postma, A. V., Gunst, Q. D., Van Den Hoff, M. J., Moorman, A. F., Wang, T. & Christoffels, V. M. 2012. Identifying the evolutionary building blocks of the cardiac conduction system. PLoS One, 7, e44231.

Keen, A. N., Fenna, A. J., Mcconnell, J. C., Sherratt, M. J., Gardner, P. & Shiels, H. A. 2016. The Dynamic Nature of Hypertrophic and Fibrotic Remodeling of the Fish Ventricle. Frontiers in Physiology, 6.

Keen, A. N., Klaiman, J. M., Shiels, H. A. & Gillis, T. E. 2017. Temperature-induced cardiac remodelling in fish. J Exp Biol, 220, 147–160.

Klaiman, J. M., Fenna, A. J., Shiels, H. A., Macri, J. & Gillis, T. E. 2011. Cardiac remodeling in fish: strategies to maintain heart function during temperature Change. PLoS One, 6, e24464.

Kochova, P., Cimrman, R., Stengl, M., Ostadal, B. & Tonar, Z. 2015. A mathematical model of the carp heart ventricle during the cardiac cycle. J Theor Biol, 373, 12–25.

Markhasin, V. S., Balakin, A. A., Katsnelson, L. B., Konovalov, P., Lookin, O. N., Protsenko, Y. & Solovyova, O. 2012. Slow force response and auto-regulation of contractility in heterogeneous myocardium. Prog Biophys Mol Biol, 110, 305–318.

Meijborg, V. M., Conrath, C. E., Opthof, T., Belterman, C. N., De Bakker, J. M. & Coronel, R. 2014. Electrocardiographic T wave and its relation with ventricular repolarization along major anatomical axes. Circ Arrhythm Electrophysiol, 7, 524–31.

Patrick, S. M., White, E., Brill, R. W. & Shiels, H. A. 2011. The effect of stimulation frequency on the transmural ventricular monophasic action potential in yellowfin tuna Thunnus albacares. J Fish Biol, 78, 651–8.

Pieperhoff, S., Bennett, W. & Farrell, A. P. 2009. The intercellular organization of the two muscular systems in the adult salmonid heart, the compact and the spongy myocardium. J Anat, 215, 536–47.

Vaĭkshnoratĭe, M. A., Belogolova, A. S., Vitiazev, V. A., Azarov, A. E. & Shmakov, D. N. 2007. [Sequence of ventricular repolarization under body cooling in the frog]. Ross Fiziol Zh Im I M Sechenova, 93, 1123–31.

Vaykshnorayte, M. A., Azarov, J. E., Tsvetkova, A. S., Vityazev, V. A., Ovechkin, A. O. & Shmakov, D. N. 2011a. The contribution of ventricular apicobasal and transmural repolarization patterns to the development of the T wave body surface potentials in frogs (Rana temporaria) and pike (Esox lucius). Comp Biochem Physiol A Mol Integr Physiol, 159, 39–45.

Vaykshnorayte, M. A., Tsvetkova, A. S. & Azarov, J. E. 2011b. Epicardial activation-to-repolarization coupling differs in the local areas and on the entire ventricular surface. J Electrocardiol, 44, 131–7.

Vornanen, M. 2016. The temperature dependence of electrical excitability in fish hearts. Journal of Experimental Biology, 219, 1941–1952.

